# Transglutaminase 2 associated with PI3K and PTEN in a putative membrane-bound signalosome platform blunts cell death

**DOI:** 10.1101/2022.02.16.480667

**Authors:** Károly Jambrovics, Pál Botó, Attila Pap, Zsolt Sarang, Zsuzsanna Kolostyák, Zsolt Czimmerer, Istvan Szatmari, László Fésüs, Iván P. Uray, Zoltán Balajthy

**Affiliations:** Department of Biochemistry and Molecular Biology, Faculty of Medicine, University of Debrecen, H-4032 Debrecen, Egyetem tér 1, Hungary; Department of Clinical Oncology, Faculty of Medicine, University of Debrecen, H-4032 Debrecen, Egyetem tér 1, Hungary

## Abstract

Atypically expressed transglutaminase 2 (TG2) has been identified as a poor prognostic factor in a variety of cancers. In this study, we evaluated the contribution of TG2 to the prolonged cell survival of differentiated acute promyelocytic leukaemia (APL) cells in response to the standard treatment with combined retinoic acid (ATRA) and arsenic trioxide (ATO). We report that one advantage of ATRA + ATO treatment compared to ATRA alone diminishes the amount of activated and non-activated CD11b/CD18 and CD11c/CD18 cell surface integrin receptors. These changes suppress ATRA-induced TG2 docking on the cytosolic part of CD18 β2-integrin subunits and reduce cell survival. In addition, TG2 overexpresses and hyperactivates the phosphatidylinositol-3-kinase (PI3K), phospho-AKT S473, and phospho-mTOR S2481 signalling axis. mTORC2 acts as a functional switch between cell survival and death by promoting the full activation of AKT. We show that TG2 presumably triggers the formation of a signalosome platform, hyperactivates downstream mTORC2-AKT signalling, which in turn phosphorylates and inhibits the activity of FOXO3, a key pro-apoptotic transcription factor. In contrast, the absence of TG2 restores basic phospho-mTOR S2481, phospho-AKT S473, PI3K, and PTEN expression and activity, thereby sensitising APL cells to ATO-induced cell death. We conclude, that atypically expressed TG2 may serve as a hub, facilitating signal transduction via signalosome formation by the CD18 subunit with both PI3K hyperactivation and PTEN inactivation through the PI3K PTEN cycle in ATRA-treated APL cells.

## INTRODUCTION

Growing evidence now supports type 2 “tissue” transglutaminase as a fundamental cell survival factor in a number of human cancers, including breast, cervical, colon, lung, and pancreatic cancers, as well as leukaemia and lymphoma^**1,2**^. TG2 can function as a signalling protein in its GTP-bound/closed/signalling□active conformations to facilitate the functions of cancer cells, but it can also work as an enzyme in its calcium-bound/open/transamidase□active form. TG2, as a multifunctional protein, shows several non-enzymatic functions that depend on its cellular location, where it plays a variety of roles in physiological and pathophysiological processes^**3**^.

The standard treatment of acute promyelocytic leukaemia (APL), a clinically and biologically distinct variant of AML, includes both targeted transcriptional and differentiation therapy and greatly induces the atypical expression of TG2 in myelocytic cells, such as NB4 cells^**4,5**^. Arsenic trioxide (ATO) has also been introduced as an APL treatment, as its addition is associated with extended survival in a large proportion of patients, even when used as a single agent^**6,7**^. However, ATO also works in synergy with ATRA to cause degradation of the PML/RARα oncoprotein^**8**^. Recent studies carried out in front-line treatment have revealed that the ATRA+ATO combination was superior to ATRA and chemotherapy, as it substantially reduced the cumulative incidences of relapse while improving disease-free and event-free survival^**9,10**^.

Using APL as a model, we have addressed the “advantages” of cancer cells expressing TG2 and now propose that the atypical expression of TG2 may alter the signal transduction pathways of these cancer cells. Here, while investigating the cell death process induced by ATO in ATRA-differentiated NB4 cell lines, we found that cell death is attenuated by TG2 expression. We discovered that the plasma membrane-bound form of TG2 functions as a signalling platform that brings CD18, c-SRC, PI3K/p110, PI3K p85, phospho-AKT T308, S473, phospho-mTOR Ser2481, and Ser2448 proteins into close proximity, thereby triggering oncogenic hyperactivity of PI3K and robust PTEN inactivation. These platforms are disintegrated into individual components by NC9 (a TG2 inhibitor) and by PP2 (a c-SRC inhibitor). TG2 therefore serves as a hub that facilitates signal transduction activation with gain-of-function PI3K and loss-of-function/tumour suppressor PTEN effects in APL.

## METHODS

### Cell lines

The following APL cell lines were cultured as described previously: NB4 WT (wildtype NB4 cells), NB4 TG2-C (virus control containing scrambled shRNA), TG2-deficient NB4 TG2-KD (shRNA-based knockdown), and NB4 TG2-KO cells (TALEN TG2 knockout)^**5,11,12**^.

### Gene expression

Methods for the isolation of RNA and RT-PCR/RT-QPCR have been published previously^**11,12**^.

### Annexin-V labelling and sorting of dead NB4 cells

1–2 × 10^6^ NB4 treated cells were harvested and labelled with FITC-conjugated Annexin-V (Biolegend) for 15 min according to the manufacturer’s instruction. Cells were analysed and sorted on a BD FACSAria III (BD Biosciences, San Jose, CA). Before sorting, first the NB4 cells were gated based on their size and the granularity, thereafter the gated and Annexin-V^+^ cells were sorted.

### Western blotting

Preparation of lysate from cell culture and WB analysis have been described previously^**11,12**^.

### Luciferase activity

Luciferase activity was measured using the Bright-Glo™ Luciferase Assay system (Promega) following the manufacturer’s instructions.

### Transfection of NB4 cells with a FOXO3A lentiviral reporter

A ready-to-transduce transcription factor-responsive lentiviral reporter system (CCS-1022L, QIAGEN) was used to generate a stable cell line, following the manufacturer’s protocol. Luciferase activity was measured as previously described^**12**^.

### Plasma membrane preparation

The plasma membranes were isolated according to the Abcam protocol (ab65400).

### PIP_3_ extraction and measurement

The isolation of PIP molecules was carried out by chloroform-methanol based organic separation. Phosphatidyl-inositol levels were determined with an ELISA detection kit (K-2500S, Echelon Biosciences) and normalized to 100 μg protein.

### *In vivo* mouse experiments

NB4 WT and NB4 TG2-KO cells (1×10^7^) were washed with sterile phosphate buffered saline (PBS) and injected into the retro-orbital region of 8–9-week-old NOD/SCID (CB17/lcr-Prkdc^scid^, JANVIER LABS) female mice under sterile conditions. Animals were divided into four treatment groups. Anaesthetised mice were administered vehicle, ATRA at 1.0 mg/kg + DEX at 50 μg, ATO at 0.75 mg/kg or combined ATRA+ATO intraperitoneally every second day for 14 days. A 27.5/0.5-gauge insulin needle/syringe (Terumo U-100, Terumo Medical Corporation, Elkton, MD) was used for the i.p. administration of treatment. DMSO administration was kept under 2% (v/v) to avoid any cytotoxic effects. At the end of the treatment, the mice were anaesthetised with isoflurane for blood sampling. Circulating human and mouse blood cells were analysed from total cardiac blood samples by flow cytometry.

All experiments were performed according to the guidelines of the Institute for Laboratory Animal Research, University of Debrecen, Faculty of Medicine, and were approved by the national and institutional ethics committee for laboratory animals used in experimental research (Project ID:4/2020/DEMAB).

### Flow cytometry for human CD11c/CD11b/Annexin-V^+^ cells

To assess the survival of human NB4 cells in mice, blood was collected and red blood cells were lysed in lysis buffer (BD PHarmn Lyse™), thereafter mouse macrophages were stained with F4/80-APC antibody (1:100x; Biolegend) for 15 min in the dark at 4°C. F4/80^-^ cells were sorted and labelled with CD11c-PE and CD11b-FITC antibodies (1:25; R&D Systems) and APC-conjugated Annexin-V (Biolegend) for 15 min at 4°C in the dark, and then analysed with a BD FACSAria III flow cytometer. The FACS data were analysed with Flowing 2.0.5 and normalised/corrected with isotype controls.

### Immunoprecipitation (IP)

IP was performed following the Thermo Fisher protocol. The isolated plasma membrane fractions were analysed by LC-MS/MS by the Proteomics Core Facility, Faculty of Medicine, University of Debrecen.

### Statistical analysis

Statistical analyses were carried out using GraphPad Prism 8.0.9 with Student’s t-test and two-way ANOVA (*P<0.05, **P<0.01, ***P<0.001).

## RESULTS

### Transglutaminase 2 lowers ATO-induced cell death in ATRA-differentiated NB4 cells

NB4 cell lines were differentiated with ATRA with or without ATO at 0.5 or 2.0 μM. Differentiation with ATRA alone stopped cell proliferation after two days in all cell lines (Figure 1A); however, the addition of ATO decreased the cell counts in a dose-dependent manner (Figure 1B, C).

**Figure 1.**
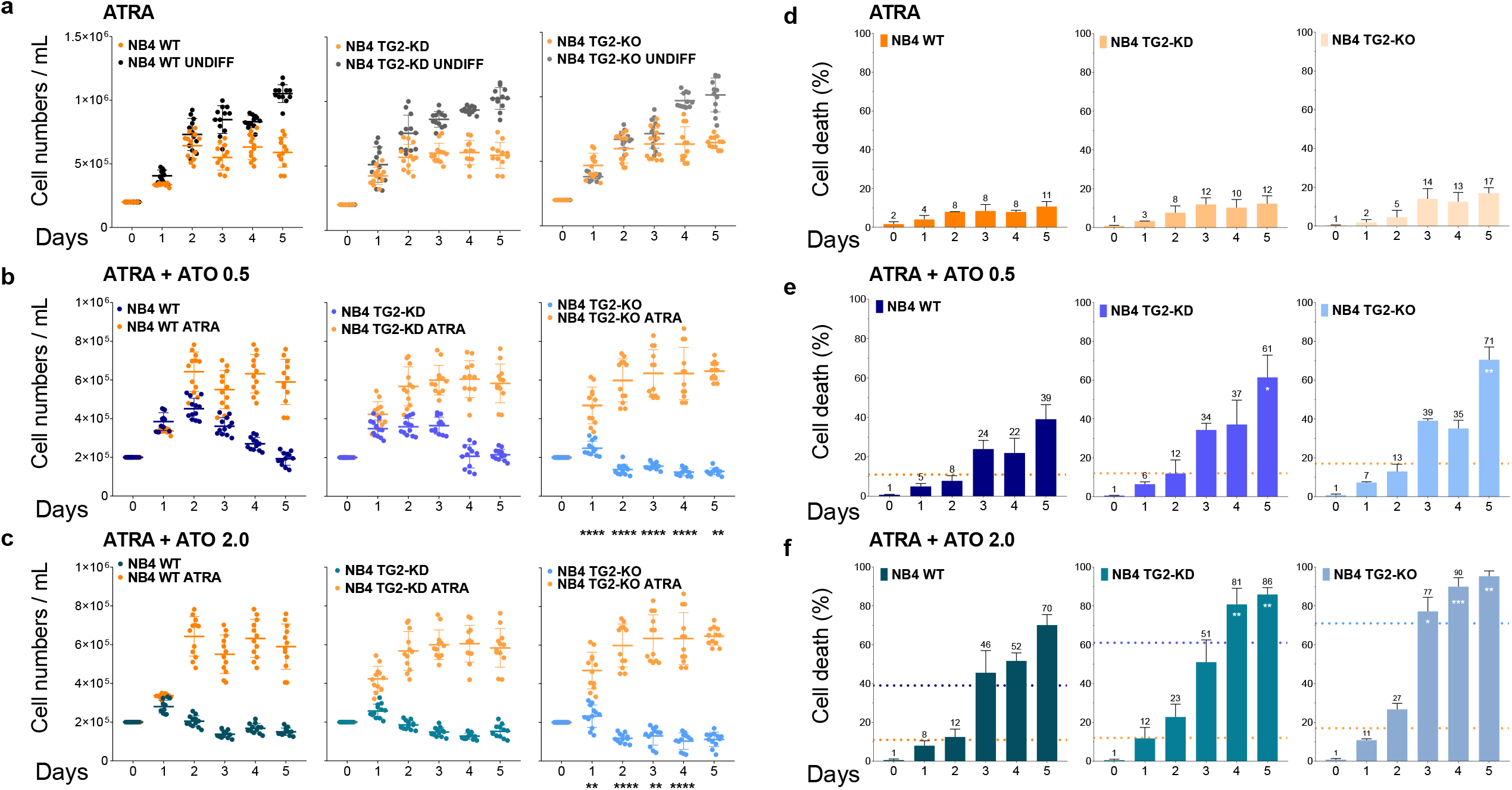
Transglutaminase 2 lowers ATO-induced cell death. (**A-C**) Cell number changes measured in KOVA Glasstic® Slide cell number counting chambers following treatment of NB4 WT, NB4 TG2-KD, and NB4 TG2-KO cells with 1 μM ATRA; 1 μM ATRA+0.5 μM ATO; and 1 μM ATRA+2.0 μM ATO at the indicated days (n=13). “Undiff” stands for undifferentiated/untreated cells. (**D-F**) FACS analysis of Annexin-V and PI stained NB4 cell lines following ATRA or ATRA+ATO treatment. The percentage of cell death ratio is represented as mean % ± SD (n=9). Measurements were conducted in triplicate; values were validated by Flowing software 2.5.1. The dotted lines intercepting the Y axis represent the mean values of ATRA (orange) or ATRA + ATO (variants of blue) treatments at day 5. Statistical analysis was conducted by two-way ANOVA (Bonferroni post-hoc test; *p <0.05, **p <0.01 and ***p <0.001, ****p <0.0001). Asterisk show the significant differences in NB4 WT vs. NB4 TG2-KO cell lines.

Annexin V-positivity in response to ATRA or ATO, determined by Flow cytometry analysis during the five-day period (Figure 1D-F), showed that ATRA induced a minor degree of cell death (Figure 1D), while TG2 induction by the ATRA+ATO (0.5 or 2.0 μM) treatment was associated with lower cell death rates in NB4 WT compared to NB4 TG2-KD cells, suggesting that cell survival was TG2-dependent (Figure 1E, day 3–5). The complete elimination of TG2 (NB4 TG2-KO) increased the susceptibility to cell death induced by ATRA+ATO treatment compared to the NB4 TG2-KD (Figure 1E day 3-5 and F, day 3-5). ATO (0.5 or 2.0 μM) treatment caused similar changes in cell numbers to ATRA + ATO (0.5 or 2.0 μM) treatment and the extent of induction of cell death is shown in Supplementary Figures 1A-F. Overall, these data indicated that TG2 expression associated with ATRA-induced differentiation contributed to cell survival.

### ATO suppresses the ATRA-induced expression of leukocyte β2 integrin receptors CD11b/CD18 and CD11c/CD18

ATRA treatment enhances the expression of CD11b/CD18 and CD11c/CD18 integrin receptors, while also activating them to a high-affinity (extended-open) form. ATO treatment significantly influences the rate of cell division and induces apoptosis. We therefore examined mRNA expression of both receptors following ATRA and ATO treatments at days 0, 3, and 5. CD11b mRNA expression showed a time-dependent, but TG2-independent, increase during ATRA treatment, whereas ATO treatment limited this expression (Figure 2A). ATO treatment alone induced a low and time-dependent mRNA expression (Supplementary Figures 2A-C). In contrast, no major differences were noted in the levels of non-active (bent-closed) CD11b/CD18 cell surface receptors in response to 0.5 or 2.0 μM ATO (Supplementary Figure 2D-E). However, on the fifth day, the expression of CD11b/CD18 receptors decreased following the combined treatments (Supplementary Figures 2M-O). The amounts of activated cell surface CD11b/CD18 receptors with a high-affinity state also increased continuously with the combined treatment (Figure 2B) but remained significantly lower than in cells treated with ATRA alone (Figure 2B).

**Figure 2.**
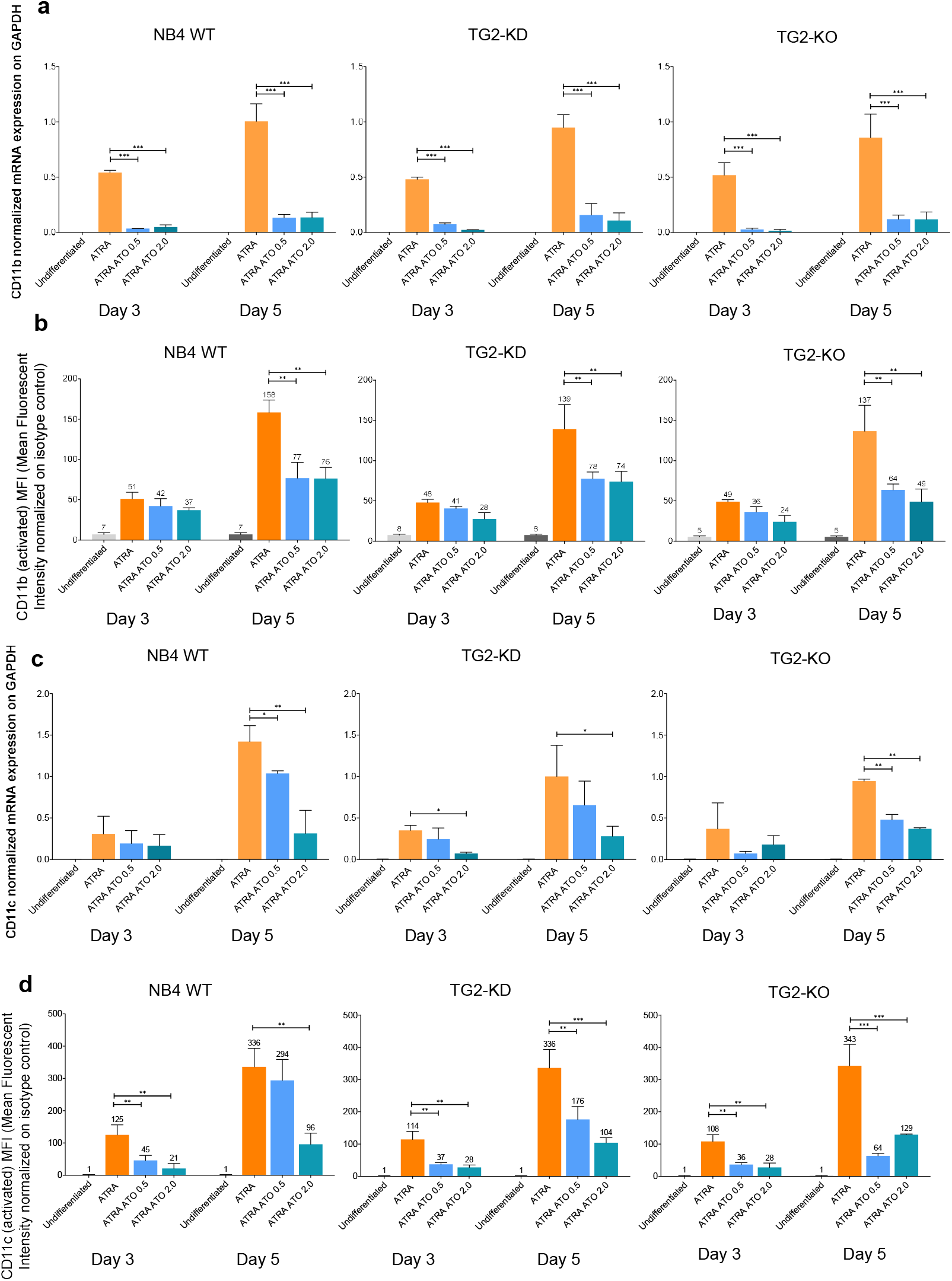
ATO suppresses the expression of leukocyte β2 integrin CD11b/CD18 and CD11c/CD18. (**A**) Relative mRNA expression of CD11b at the indicated days by RT-qPCR normalized to GAPDH (n=3). (**B**) Flow cytometry analysis of cell surface expression of the differentiation marker CD11b/CD18 (n=3). (**C**) Relative mRNA expression of CD11c measured by RT-qPCR normalized to GAPDH (n=3). (**D**) Flow cytometry analysis of cell surface expression of differentiation marker CD11c/CD18 (n=3). Measurements were conducted in triplicate; values were validated by Flowing software 2.5.1. Statistical analysis was conducted by two-way ANOVA (Bonferroni post hoc test; *p <0.05, **p <0.01 and ***p <0.001, ****p <0.0001).

CD11c mRNA expression was barely induced by single ATO treatments in NB4 cell lines (Supplementary Figures 2G-I). In contrast, the CD11c mRNA levels increased robustly and in a time-dependent manner with ATRA treatment, but this increase was mitigated in a concentration-dependent manner by the addition of ATO (Figure 2C). Comparison of the expression of non-active CD11c/CD18 (bent-closed) cell surface receptors revealed that the addition of ATO blunted the ATRA-induced CD11c/CD18 expression (Supplementary Figures 2P-R). The use of an antibody (Clone 3.9) that detected the activated (extended-open) form of cell surface CD11c/CD18 revealed a similar response to ATRA and ATO treatments to that seen for mRNA (Figure 2D). Thus, these data indicate that the activated and non-activated forms of CD11b/CD18 and CD11c/CD18 receptors remained significantly lower with the combined treatment than in cells treated with ATRA alone.

### ATRA-induced TG2 boosts the phosphorylation of mTORC2 and keeps FOXO3 transcriptionally inactive

The PI3K, AKT/Protein Kinase B, and mTOR signalling pathways are critical for the regulation of cell growth and survival. mTORC1 is phosphorylated mainly on Ser2448 of mTOR, whereas mTORC2 is mainly phosphorylated on Ser2481 of mTOR^**13,14**^. ATRA-differentiated NB4 cells showed phospho-S2481-mTOR levels that changed with the amount of TG2. The appearance of phospho-S2481-mTORC was significantly higher in differentiated WT NB4 and virus control NB4 C cells than in NB4 TG2-KD or TG2-deficient NB4 TG2-KO cells, and it was also associated with the decreased phosphorylation of AKT at Ser473 (Figure 3 A, B and C, D; first block). The degree of phosphorylation of mTOR S2448 did not correlate with TG2 expression (Figure 3A, first block).

**Figure 3.**
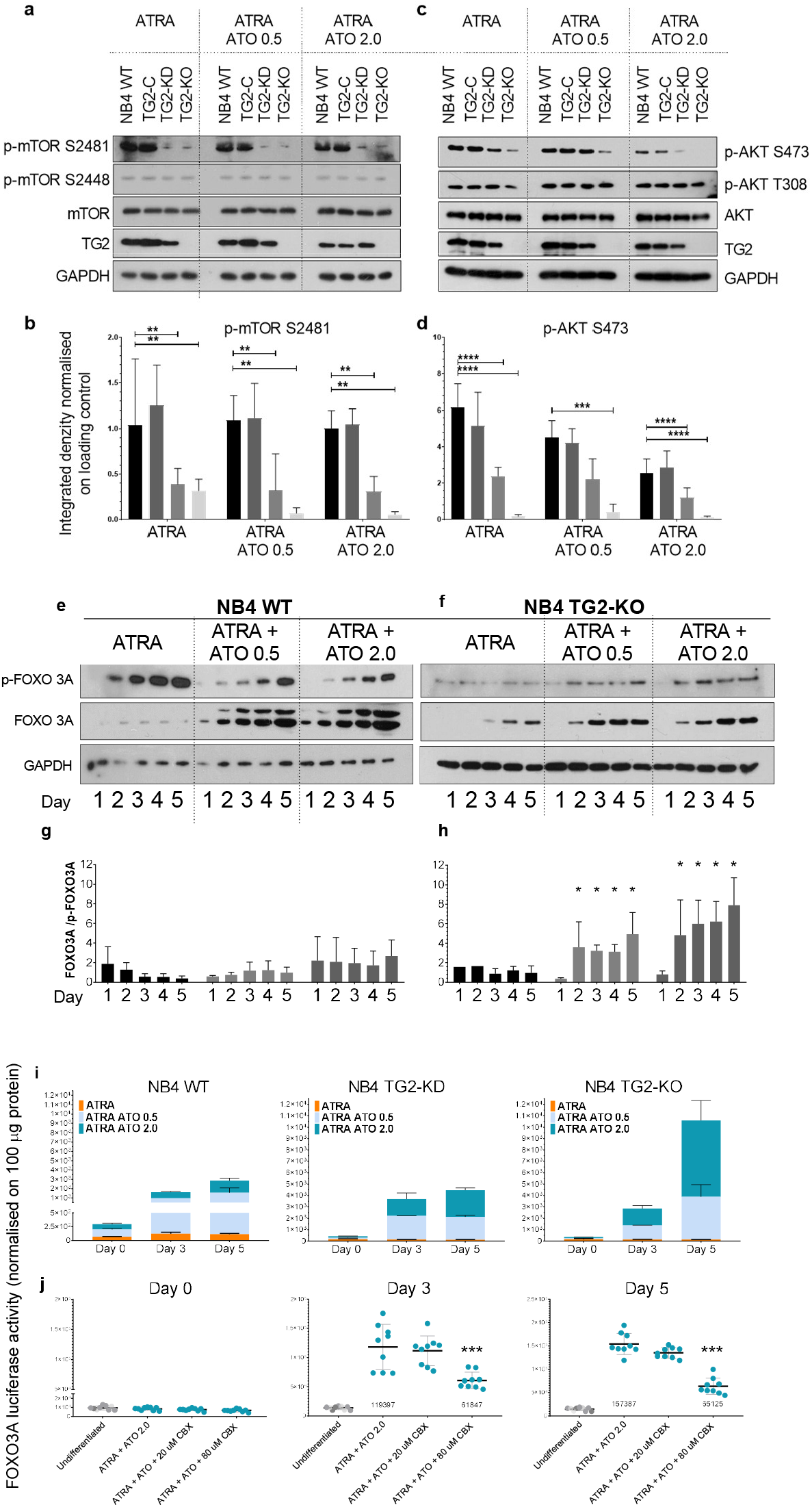
TG2 boosts the phosphorylation of mTORC2 and maintains FOXO3 transcriptionally inactive. **(A, C)** Representative western blot showing p-mTOR (2481, 2448) p-AKT (S473, T308), mTOR, AKT, TG2, and GAPDH protein expression levels in NB4 cell lines after 5 days (n=9). (**B, D**) Densitometry analysis results represent integrated optical density normalized to a loading control. (**E, F**) Representative western blot showing p-FOXO1/FOXO3/FOXO4 (T24/T32), FOXO3A and GAPDH protein expression levels for 5 days (n=5). (**G, H**) Densitometric analysis of the FOXO3A and phosphorylated FOXO3A blots. (**I**) NB4 cell lines containing a FOXO3A luciferase reporter element were treated and measured by a luminescence-based method in triplicate and reported as relative light units (RLU) (n=3). Values are normalized to 100 μg cell lysate protein from each cell line. (**J**) NB4 cell lines containing the FOXO3A luciferase reporter element were treated with ATRA or ATRA+ATO with or without the FOXO3A inhibitor CBX and measured by the luminescence-based method in triplicate and reported as RLU. Values are normalized to 100 μg cell lysate protein from each cell line. Statistical analysis was conducted by two-way ANOVA (Bonferroni post hoc test; *P<0.05, **P<0.01 and ***P<0.001, ****P<0.0001).

PIP_3_ activates PDK1 to trigger the phosphorylation of AKT at Thr308^**21**^. Although AKT Thr308 phosphorylation appeared to be TG2-independent, the 473-serine side chain was phosphorylated to a significantly greater extent at ATRA-induced TG2 protein levels (in NB4 WT and NB4 TG2-C cells) than when TG2 expression was low or absent (in NB4 TG2-KD and NB4 TG-KO cells) (Figure 3 C, D, first block). The combination of ATRA with either 0.5 or 2.0 μM ATO had the same effect on the amount of phospho-S2481-mTOR as ATRA alone (Figure 3 A, B, second and third blocks). By contrast, a concentration-dependent suppression of AKT Ser473 phosphorylation occurred after ATO treatments and was independent of the levels of S2481-phosphorylation of mTOR (Figure 3 C, D, second and third blocks).

FoxO transcription factors are major downstream targets of the PI3K-AKT signalling pathway controlling cell proliferation and survival. Phosphorylation by AKT results in the nuclear export and cytoplasmic sequestration of FOXO, thereby inhibiting its transcriptional activity and promoting cell survival, growth, and proliferation^**22,23,24**^. In ATRA-treated cells, phosphorylated-FOXO3 amounts increased gradually relative to total FOXO3 expression until day 5 (Figure 3E, first block). In contrast, ATRA+ATO treatment attenuated and delayed the induction of phosphorylated FOXO3, while significantly increasing the total amount of FOXO3 protein (Figure 3 E, second and third blocks). In the absence of TG2, the phosphorylated FOXO3 levels were practically unchanged in ATRA-treated NB4 TG2-KO cells up to day 5, whereas the total FOXO3 protein levels increased slightly from day 3 to day 5, ATRA+ATO treatment intensified that (Figure 3F, H).

The examination of FOXO3 transcriptional activity using the FOXO3A promoter-driven luciferase construct stably integrated into the NB4 cell genome revealed that increasing ATO concentrations were associated with enhanced luciferase reporter activity in cells expressing no or low TG2 levels (Figure 3I). TG2 expression inhibited the transcriptional activity of FOXO3 (Figure 3I, first column) via the AKT S473 phosphorylated form (Figure 3C and D, first block). In the presence of ATO and the absence of TG2, AKT S473 remained unphosphorylated (Figure 3C and D, third blocks) and was associated with a significant increase in FOXO3 transcriptional activity (Figure 3I, third column). Treatment with carbenoxolone (CBX), a potential FOXO3 inhibitor, reduced the luciferase reporter activity driven by the FOXO3A promoter by 48.2% at day 3 and 58.7% at day 5 in ATRA+ATO treated NB4 WT cells (Figure 3J). Annexin-V positivity decreased by 13% and 9% at days 3 and 5, respectively, in parallel with the reduced FOXO3 transcriptional activity (Supplementary Figure 3). Collectively, these data indicate a potential role of TG2 to increase phosphorylation of mTOR on Ser2481 suppressing transcriptional activity of FOXO3.

### TG2 is required for enhanced PI3K-AKT-TOR signal transduction pathway activity

Our working hypothesis was that atypical TG2 expression sustains cell survival by activating the mTORC2, Phospho-AKT (S473, T308), and Phospho-FOXO3 signalling pathways. The binding of growth factors to a RTK can activate PI3K to generate PIP_3_ whose concentration has been determined in NB4 cell lines. The PIP_3_ results confirmed that ATRA treatment of NB4 WT cells increased the PIP_3_ level by about 180,000-fold compared to NB4 TG2-KO cells. The PIP_3_ levels were 11,000-fold higher in NB4 TG2-KD cells than in TG2-KO cells (Figure 4A), and this trend was not affected by the ATRA+ATO treatment (Figure 4B). The PIP_3_ content of undifferentiated dividing NB4 cell lines was about one-tenth that of differentiated non-dividing NB4 WT cells (Supplementary Figure 4). Sorting of the ATRA- and ATRA+ATO-treated NB4 cell lines for live and apoptotic cells revealed PIP_3_ concentrations that were 7-fold higher in Annexin-V negative NB4 WT cells and 2-fold higher in TG2-KD cells than in NB4 TG2-KO cells (Figure 4C). The PIP_3_ levels were significantly lower in apoptotic Annexin-V-positive cells of every cell line compared to live cells (Figure 4D).

**Figure 4.**
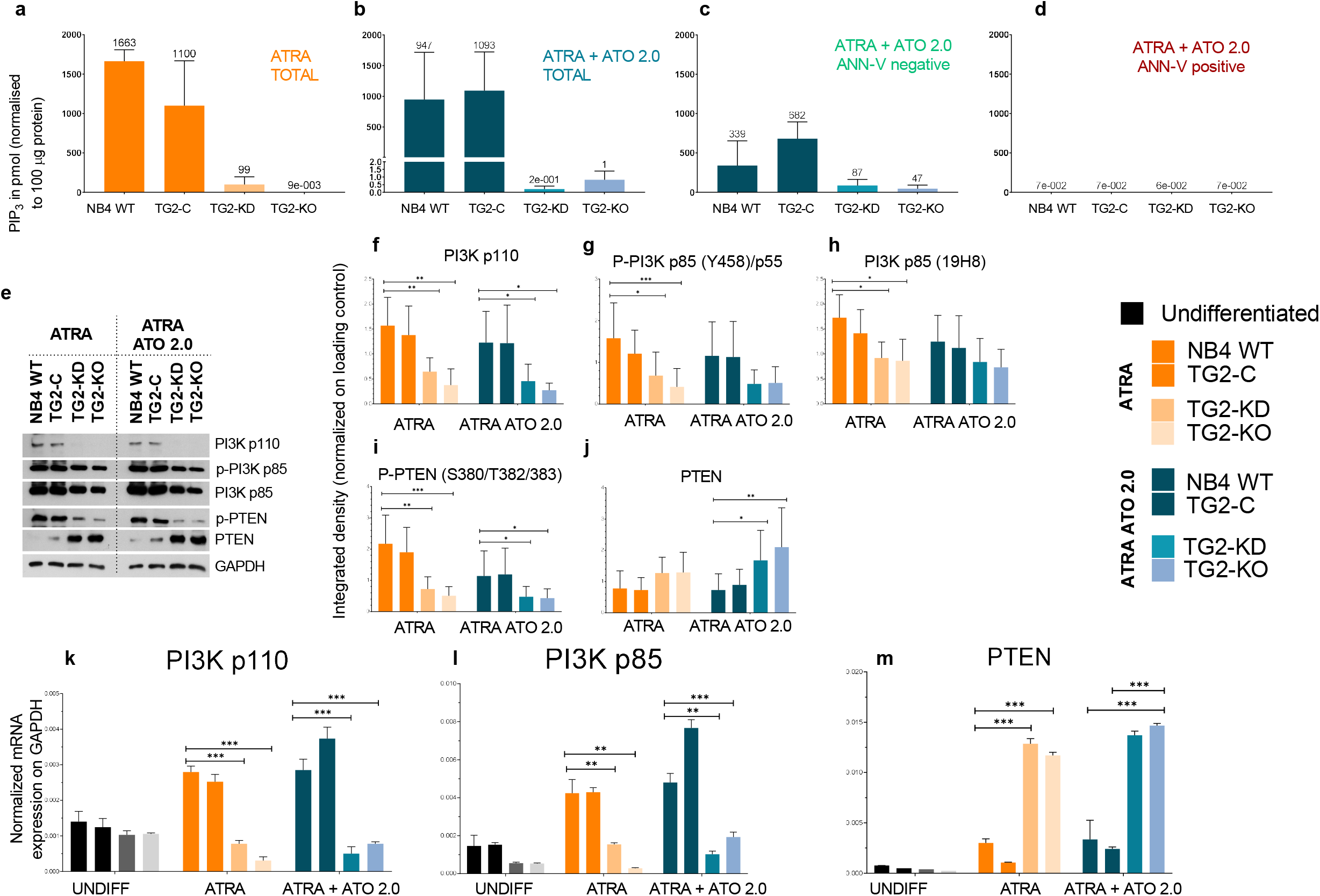
TG2 is required for enhanced PI3K-AKT-TOR signal transduction pathway activity. **(A-D)** Membrane PIP_3_ levels were quantified by an ELISA-based fluorescence detection method (Echelon Inc.). Figures show the representation of the mean values ± SD of PIP_3_ levels from three independent experiments (n=3). Measurements were conducted in triplicate. (**A**) PIP_3_ levels in ATRA-treated total NB4 cells (**B**) PIP_3_ levels in ATRA+ATO treated unsorted NB4 cells (**C**) PIP_3_ levels in ATRA+ATO-treated Annexin-V labeled but Annexin-V negative cells (**D)** PIP_3_ levels in ATRA+ATO-treated Annexin-V labeled and Annexin-V positive cells (n=3) (**E**) Representative western blot showing PI3K subunits, p-PTEN, PTEN, and GAPDH protein expression levels in NB4 cells treated for 5 days with ATRA or ATRA+ATO (n=5). (**F-J**) Densitometry analysis of western blots of PI3K pathway proteins. (**K-M**) Relative mRNA expression of *PI3K*-*p110, PI3K-p85*, and *PTEN* in NB4 cells treated with 1 μM ATRA or 1 μM ATRA+ATO; RT-qPCR results are normalized to GAPDH (n=3). Statistical analysis was conducted by two-way ANOVA (Bonferroni post hoc test; *p <0.05, **p <0.01 and ***p <0.001, ****p <0.0001).

The efflux of PIP_3_ is determined by the amounts of monomeric PI3K/p110, PTEN, and phospho-PTEN (Figure 4E). The expression of PI3K/p110, the PI3K p85 regulatory subunit, and phospho-PI3K p85 showed TG2 dependency (Figure 4F-H, first four columns). High TG2 expression was accompanied by increased amounts of PI3K/p110 (Figure 4F), phospho-PI3K p85 (Figure 4G), PI3K p85 (Figure 4H), and phospho-PTEN (Figure 4I), whereas low (TG2-KD) or no (TG2-KO) TG2 expression was associated with levels one-third and one-fifth of WT, respectively. The phosphatase-active PTEN expression showed a moderate inverse correlation with TG2 expression (Figure 4J, first four columns), whereas the phosphatase-inactive p-PTEN displayed a powerful proportional correlation with TG2 (Figure 4I, first four columns). Phosphatase-inactive phospho-PTEN patterns also correlated with patterns of PI3K/p110 and phospho-PI3K p85 expression, suggesting that TG2 induction may support PIP_3_ synthesis (Figure 4F, G, and I, first two columns), whereas low TG2 expression was associated with minimal PIP_3_ synthesis (Figure 4F, G, and I; third and fourth columns). In WT cells, ATRA+ATO treatment moderately decreased the amount of phosphatase-inactive phospho-PTEN (Figure 4I, right block), whereas phosphatase-active PTEN was increased in TG2 KD and KO cells (Figure 4J, right block). PI3K, PI3K p85 and PTEN mRNA expressions were strongly correlated with the Western-blot results shown in Figure 4 K-M. The levels of PI3K p110 mRNA, PI3K p85 mRNA, and PTEN mRNA (Figure 4K-M) showed apparent TG2 dependence. Together, our data show that, while apoptotic cells contain virtually no PIP3, differentiated NB4 WT cells expressing TG2 possess ten times more PIP3 than dividing undifferentiated cells expressing no TG2.

### TG2 sustains cell survival by forming a signalosome platform and hyperactivating the PI3K p-AKT S473 and p-mTORC2 signalling axis

The *in vivo* requirement for TG2 for the survival of NB4 cells was examined by the intravenous injection of 1×10^7^ NB4 WT and TG2-KO cells via the suborbital vein into NOD/SCID mice^**25**^. After 14 days, FACS and Alu-based real-time PCR methods confirmed that blood samples from control, ATRA-treated, and ATRA+ATO-treated mice contained comparable numbers of NB4 WT and TG2-KO cells^**11,20**^. Only the ATRA+ATO treatment reduced the NB4 TG2-KO cell counts below WT levels (Figure 5A).

**Figure 5.**
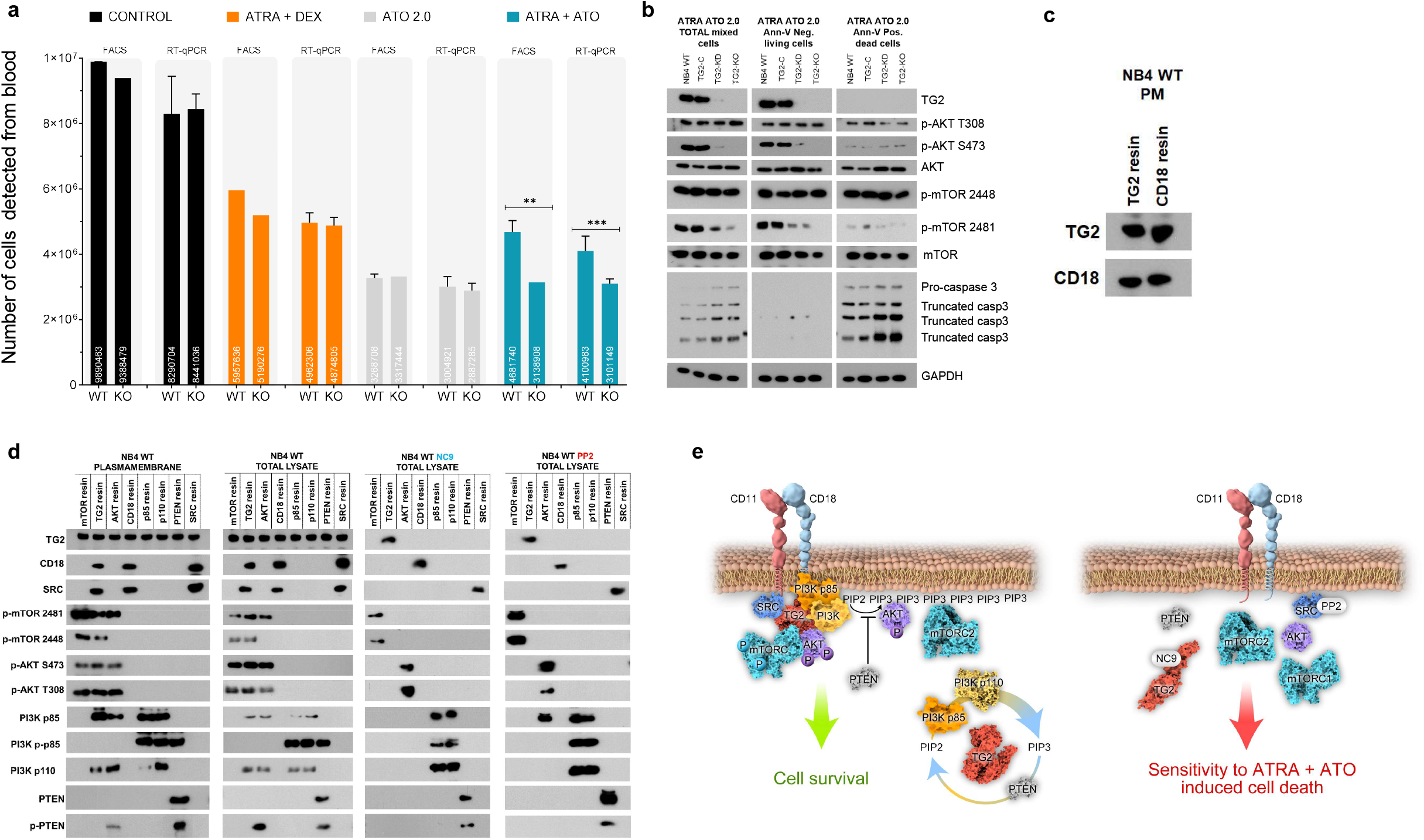
TG2 sustains cell survival by forming a signalosome platform and hyperactivating the PI3K p-AKT S473 and p-mTORC2 signaling axes. **(A)** NB4 WT and TG2-KO cells (1×10^7^) were intravenously injected via the suborbital vein into NOD/SCID female mice, followed by intraperitoneal injection of ATRA 1.0 mg/kg + DEX at 50 μg, ATO at 0.75 mg/kg, or ATRA+ATO every two days. At day 14, blood samples were collected from the heart and analyzed for total numbers of NB4 WT and TG2-KO cells by FACS and human-ALU-based RT-qPCR analyses. Graphs show the mean ± SD values after 14 days (n=4). Statistical analysis was conducted by two-way ANOVA (Bonferroni post hoc test; *p <0.05, **p <0.01 and ***p <0.001, ****p <0.0001). (**B**) Representative western blot showing expression of TG2, p-AKT (S473, T308), AKT, p-mTOR (2481, 2448), mTOR, pro-caspase3/truncated forms, and GAPDH proteins in the total cell lysate or in a cell population sorted by Annexin-V labeling (n=7). (**C**) Representative IP-western blot showing plasma membrane CD18 and TG2 protein expression levels following ATRA treatment of NB4 WT cells (n=5). (**D**) Representative co-immunoprecipitation-Western-blot showing plasma membrane and total cell lysate TG2, CD18, SRC, p-mTOR (2481, 2448), p-AKT (S473, T308), p85, p-p85, p110, PTEN, and p-PTEN protein expression levels in NB4 WT cells following upon ATRA, ATRA+ NC9 (30 μM), or ATRA+PP2 (20 μM) treatment (n=3). (**E**) TG2 - dependent signalosome formation and its inhibition by NC9 or PP2.

The antiapoptotic effect of TG2 was then investigated by sorting the ATRA+ATO-treated NB4 cells by annexin-V staining and analysing them by Western blotting for the expression of TG2, AKT, phospho-AKT (S473, T308), mTOR, phospho-mTOR (S2448, S2481), and procaspase 3 and its truncated forms (Figure 5B). The immunoblotting results showed reduced amounts of AKT S473 phosphorylation at low TG2 expression, with both being undetectable in TG2-KO cells. Phospho-mTOR S2481 expression also showed TG2 dependence. Apoptotic markers, in the form of cleaved caspase 3 and procaspase 3, were present in higher amounts in cells with lower TG2 expression (e.g., NB4 TG2-KD or TG2-deficient TG2-KO cells) (Figure 5B, left block). Significant amounts of TG2 were detected in the living Annexin-V-negative WT NB4 and TG2-C cells, but not in the TG2-KD cells. The phosphorylation of AKT S473 and mTOR 2481 decreased in proportion to the amount of TG2. Procaspase 3 and its cleaved forms were below the detection limit (Figure 5B, middle panel).

In Annexin-V-positive apoptotic NB4 WT, TG2-C, and TG2-KD cells, no TG2 was detected, and AKT T308 phosphorylation was reduced in all cell lines. However, the loss of TG2 in dying cells was associated with the suppression of phospho-AKT S473 and phospho-mTOR S2481 levels. The levels of procaspase 3 and its cleavage forms were clearly higher than in the unsorted cell populations (Figure 5B, right panel).

The synthesis of PIP_3_ was localised to the cell membrane and appeared to depend on TG2 (Figure 4A-M), prompting the question of how TG2, which is commonly considered a cytoplasmic protein, might form an association with the plasma membrane. Mass spectrometry analysis of the plasma membrane fraction of NB4 cells identified several hundred proteins. Of these, the CD18 integrin protein was pulled down in a complex that was immunoprecipitated with a TG2 antibody. TG2 also co-immunoprecipitated with CD18; therefore, we concluded that CD18 forms plasma membrane–associated protein complexes with TG2 in differentiated NB4 cells (Figure 5C). TG2 was detected in a membrane-associated form, leading us to suspect that TG2 might cooperate with proteins, such as c-SRC, PI3K/110, PTEN, and mTOR, that lie downstream in the PI3K-AKT-mTOR signal transduction cascade and participate in protein-protein interactions. Co-immunoprecipitation experiments with the polyclonal TG2 antibody detected interactions between the TG2, CD18, c-SRC, phosphorylated mTOR (S2481, S2448) from mTORC1 and mTORC2, phospho-AKT (S473, T308), PI3K/p110 and PI3K phospho-p85 (Figure 5D, 1st and 2nd vertical panel). At the same time, antibodies to mTOR, AKT, CD18, p85, PI3K/p110, PTEN, and c-SRC also co-immunoprecipitated with TG2 in the reciprocal co-immunoprecipitation experiments (Figure 5D, 1st and 2nd horizontal panel). Treatment with either the NC9 TG2 inhibitor or the PP2 c-SRC inhibitor abolished all interactions between TG2 and its interacting proteins. These experiments showed that TG2 supports cell survival *in vivo* based on the formerly unknown capability of TG2 to form signalosome platforms by attracting CD18.

## Discussion

The normal front-line therapy for APL has been the simultaneous administration of ATRA and chemotherapy. However, clinical studies have now revealed that ATO, either alone or in combination with ATRA, can improve the outcomes of APL. Here, we showed that the survival of APL leukaemic cells depends on an active PI3K-AKT-mTORC pathway and a transcriptionally inactive FOXO3 signalling axis TG2 dependent manner in the combined treatments.

TG2 is known to modulate gene expression, and our findings show that it can reciprocally regulate *PI3K* and *PTEN*. However, the differences in enzyme activities in the presence or absence of TG2 expression does not explain the nearly 180,000-fold difference in PIP_3_ levels, suggesting that TG2 also enhances PIP_3_ production through some other mechanism that does not involve gene expression differences. PIP_3_ is present in minute amounts in the membranes of apoptotic cells (Figure 4D), suggesting that the PI3K-AKT survival signalling pathway is inactivated in dying cells, whereas live Annexin-V negative cells contain high PIP_3_ levels (Figure 4C). Our examination of TG2 in Annexin-V negative live and Annexin-V positive apoptotic cells showed that the amount of TG2 is below the detection limit in Annexin-V-positive apoptotic cells but is measurable in Annexin-V-negative cells (Figure 5B 2nd and 3rd panels).

TG2 in its GTP-bound/closed/signalling□active form acts as an organisational constituent or a signalosome for PI3K-AKT-mTOR-FOXO3 signal transduction by promoting the assembly of multiprotein signal transduction complexes at the plasma membrane. This enables the establishment of false spatially concentrated and site-specific signals with atypically increased PI3K, phospho-mTOR S2481, and phospho-AKT S473 signalling activities. Both the specificity and the regulation of signal transduction are extraordinarily increased, since certain signalling proteins, such as CD18, c-SRC, phospho-AKT T308, S473, phospho-mTOR S2481 and S2448, can associate with the GTP-bound/closed/signalling□active TG2. The open/transamidase□active conformation is locked by NC9, a TG2-selective irreversible inhibitor, thereby terminating GTP binding and the TG2 signalosome. Treatment with PP2, an inhibitor of SRC, can also accomplish this.

In summary, the atypical expression of GTP-bound TG2 is an upstream signal activating a signalosome platform that hyperactivates a PI3K, phospho-AKT S473, and phospho-mTOR S2481 signalling axis via CD18, along with a robust PTEN inactivation, to maintain phospho-FOXO3 in a transcriptionally inactive state and promote cell survival. TG2 is dysregulated in many tumours; therefore, reducing TG2 levels and/or modifying the open/transamidase□active form may represent promising strategies to sensitise tumour cells to cancer therapy.

## Acknowledgement

The authors would like to thank Jeffrey W. Keillor for the NC9 compound and Tímea Silye-Cseh for assistance with mice experiments.

## References

1. Eckert RL, Fisher ML, Grun D, Adhikary G, Xu W, Kerr C. Transglutaminase is a tumor cell and cancer stem cell survival factor. Molecular carcinogenesis. 2015;54(10):947–58.

2. Chen F, Zhang Y, Varambally S, Creighton CJ. Molecular correlates of metastasis by systematic pan-cancer analysis across The Cancer Genome Atlas. Molecular Cancer Research. 2019;17(2):476–87.

3. Tabolacci C, De Martino A, Mischiati C, Feriotto G, Beninati S. The role of tissue transglutaminase in cancer cell initiation, survival and progression. Medical Sciences. 2019;7(2):19.

4. Benedetti L, Grignani F, Scicchitano BM, Jetten AM, Diverio D, Lo Coco F, et al. Retinoid-induced differentiation of acute promyelocytic leukemia involves PML-RARalpha-mediated increase of type II transglutaminase. 1996.

5. Balajthy Z, Csomós K, Vámosi G, Szántó A, Lanotte M, Fésüs L. Tissue-transglutaminase contributes to neutrophil granulocyte differentiation and functions. Blood. 2006;108(6):2045–54.

6. Sanz MA, Lo-Coco F. Modern approaches to treating acute promyelocytic leukemia. Journal of Clinical Oncology. 2011;29(5):495–503

7. Sanz MA, Montesinos P. How we prevent and treat differentiation syndrome in patients with acute promyelocytic leukemia. Blood. 2014;123(18):2777–82.

8. de Thé H, Pandolfi PP, Chen Z. Acute promyelocytic leukemia: a paradigm for oncoprotein-targeted cure. Cancer Cell. 2017;32(5):552–60.

9. Zhu H-H, Hu J, Lo-Coco F, Jin J. The simpler, the better: oral arsenic for acute promyelocytic leukemia. Blood. 2019;134(7):597–605.

10. Cicconi L, Platzbecker U, Avvisati G, Paoloni F, Thiede C, Vignetti M, et al. Long-term results of all-trans retinoic acid and arsenic trioxide in non-high-risk acute promyelocytic leukemia: update of the APL0406 Italian-German randomized trial. Leukemia. 2020;34(3):914–8.

11. Csomós K, Német I, Fésüs L, Balajthy Z. Tissue transglutaminase contributes to the all-trans-retinoic acid–induced differentiation syndrome phenotype in the NB4 model of acute promyelocytic leukemia. Blood. 2010;116(19):3933–43.

12. Jambrovics K, Uray IP, Keresztessy Z, Keillor JW, Fésüs L, Balajthy Z. Transglutaminase 2 programs differentiating acute promyelocytic leukemia cells in all-trans retinoic acid treatment to inflammatory stage through NF-κB activation. haematologica. 2019;104(3):505–15.

13. Mirabilii S, Ricciardi MR, Piedimonte M, Gianfelici V, Bianchi MP, Tafuri A. Biological Aspects of mTOR in Leukemia. International Journal of Molecular Sciences. 2018;19(8):2396.

14. Liu P, Gan W, Chin YR, Ogura K, Guo J, Zhang J, et al. PtdIns (3, 4, 5) P3-dependent activation of the mTORC2 kinase complex. Cancer discovery. 2015;5(11):1194–209.

15. Sarbassov DD, Guertin DA, Ali SM, Sabatini DM. Phosphorylation and regulation of Akt/PKB by the rictor-mTOR complex. Science. 2005;307(5712):1098–101.

16. Eijkelenboom A, Burgering BM. FOXOs: signalling integrators for homeostasis maintenance. Nature reviews Molecular cell biology. 2013;14(2):83–97.

17. Zhang X, Tang N, Hadden TJ, Rishi AK. Akt, FoxO and regulation of apoptosis. Biochimica et Biophysica Acta (BBA)-Molecular Cell Research. 2011;1813(11):1978–86.

18. Liu Y, Ao X, Ding W, Ponnusamy M, Wu W, Hao X, et al. Critical role of FOXO3a in carcinogenesis. Molecular cancer. 2018;17(1):1–12.

19. Wang H-Y, Zhang B, Zhou J-N, Wang D-X, Xu Y-C, Zeng Q, et al. Arsenic trioxide inhibits liver cancer stem cells and metastasis by targeting SRF/MCM7 complex. Cell death & disease. 2019;10(6):1–16.

20. Funakoshi K, Bagheri M, Zhou M, Suzuki R, Abe H, Akashi H. Highly sensitive and specific Alu-based quantification of human cells among rodent cells. Scientific reports. 2017;7(1):1–12

