## Supplement for "Transglutaminase 2 associated with PI3K and PTEN in a putative membrane-bound signalosome platform blunts cell death"

#### Supplement Figure 1.

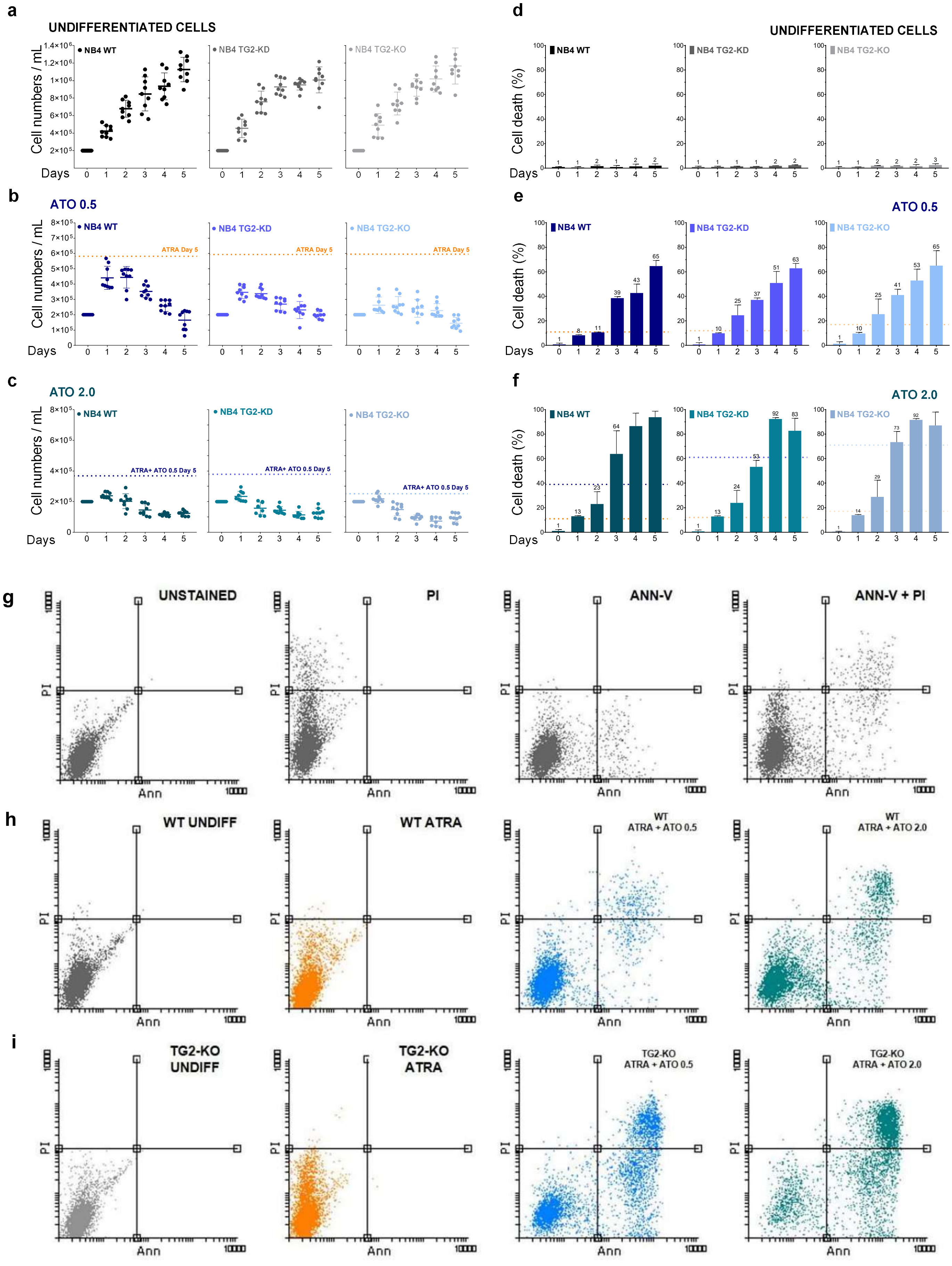

### Supplemental Figure 2.

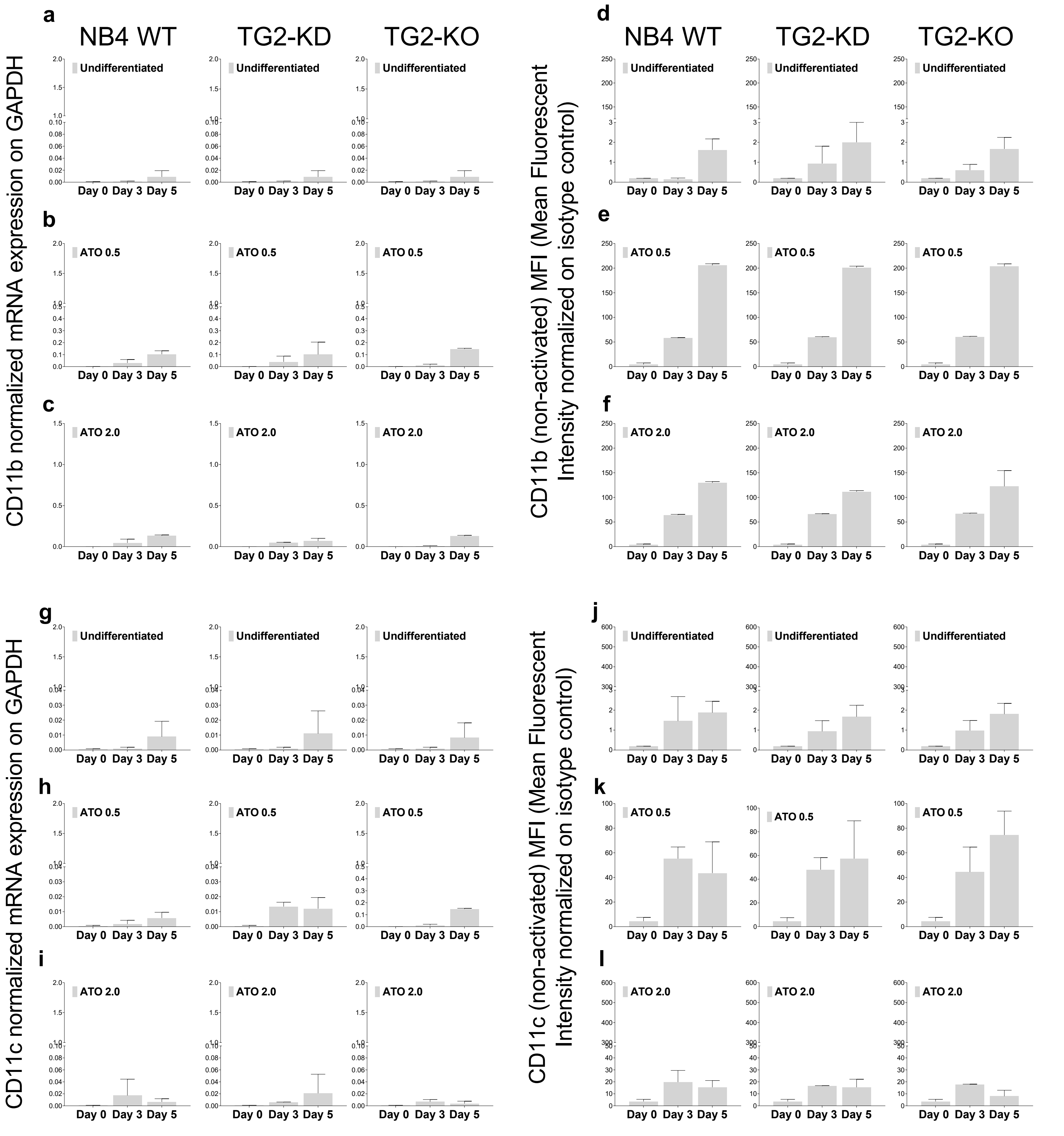

#### Supplemental Figure 2.

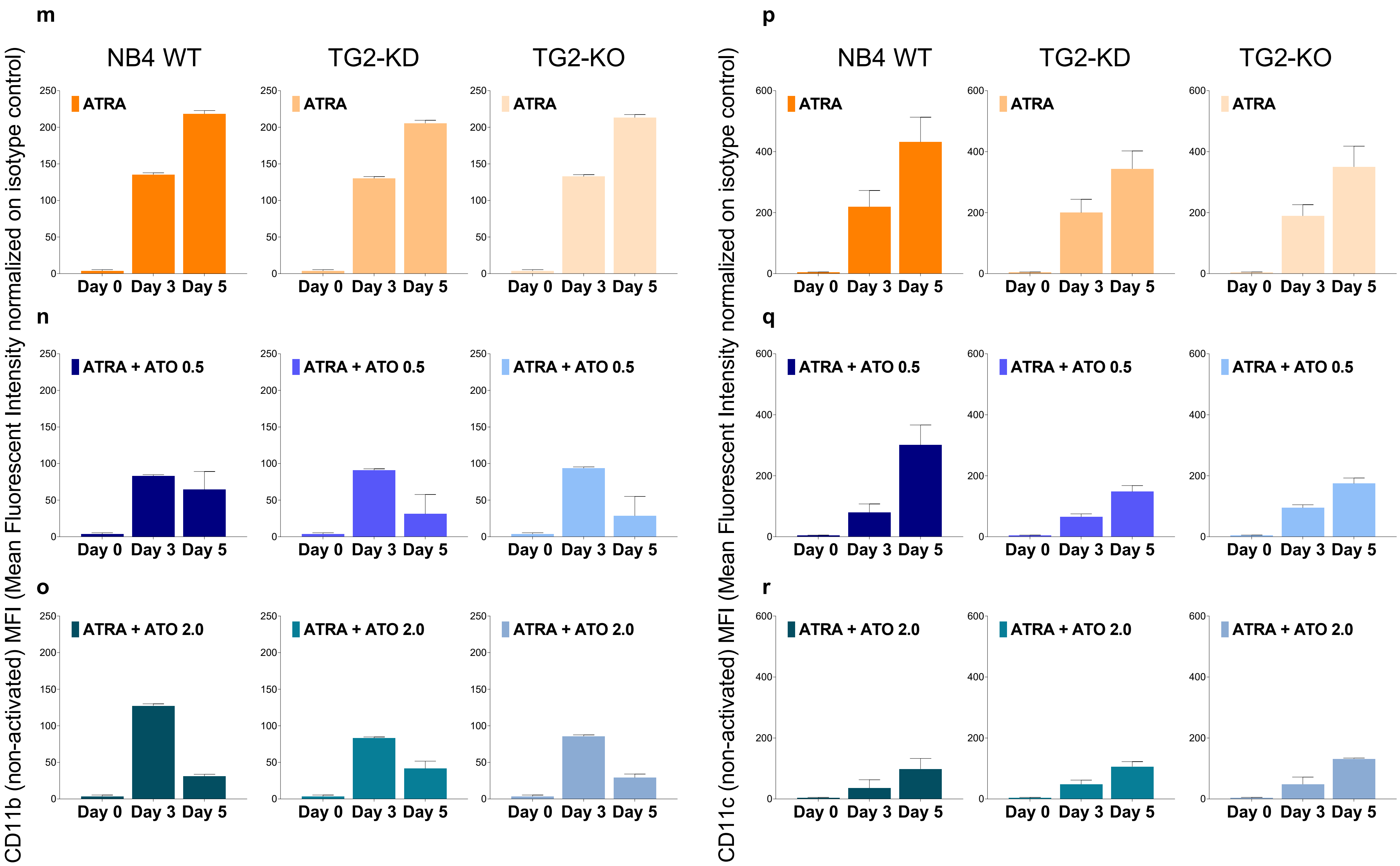

**Supplement Figure 3.**

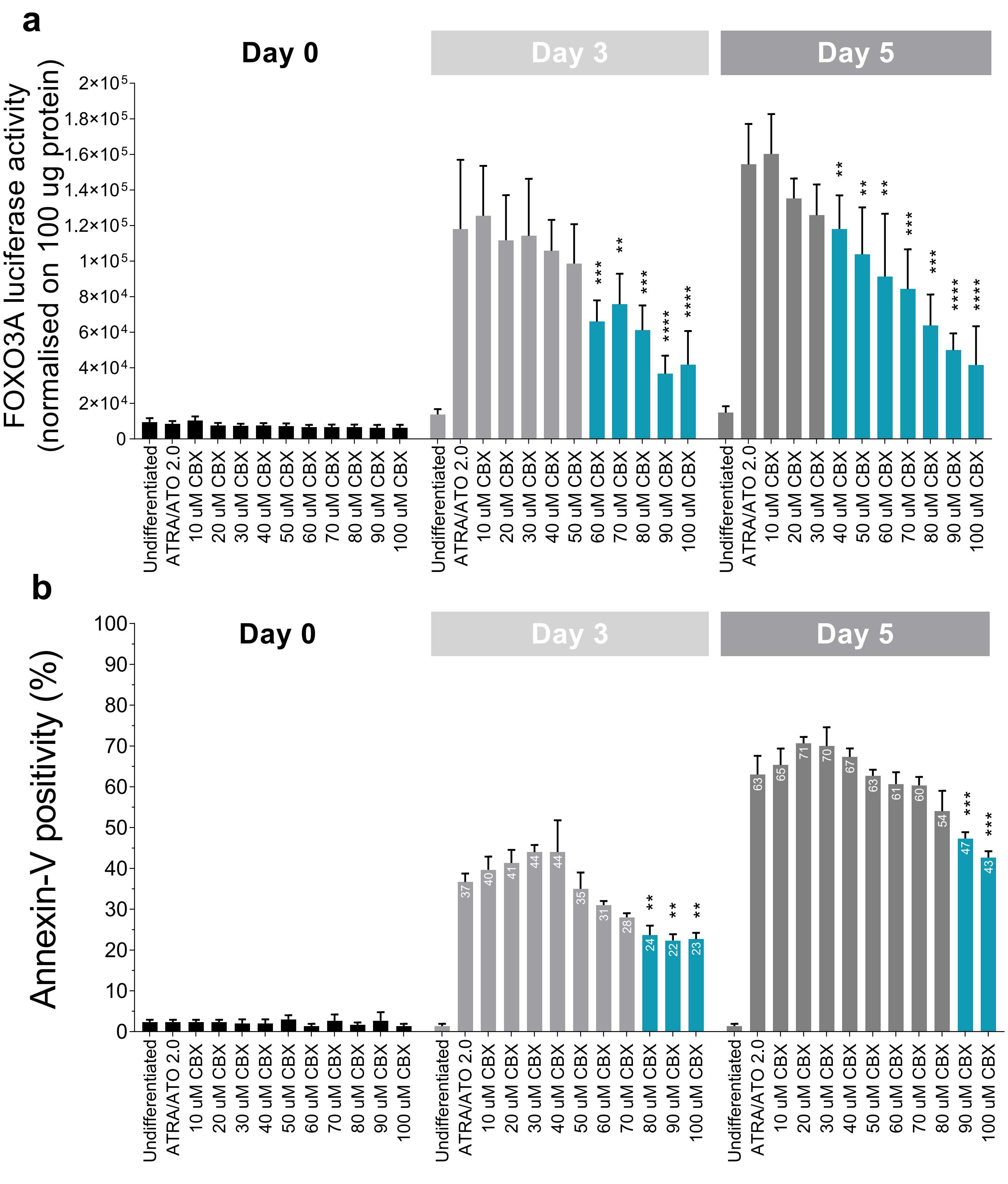

**Supplemental Figure 4.**

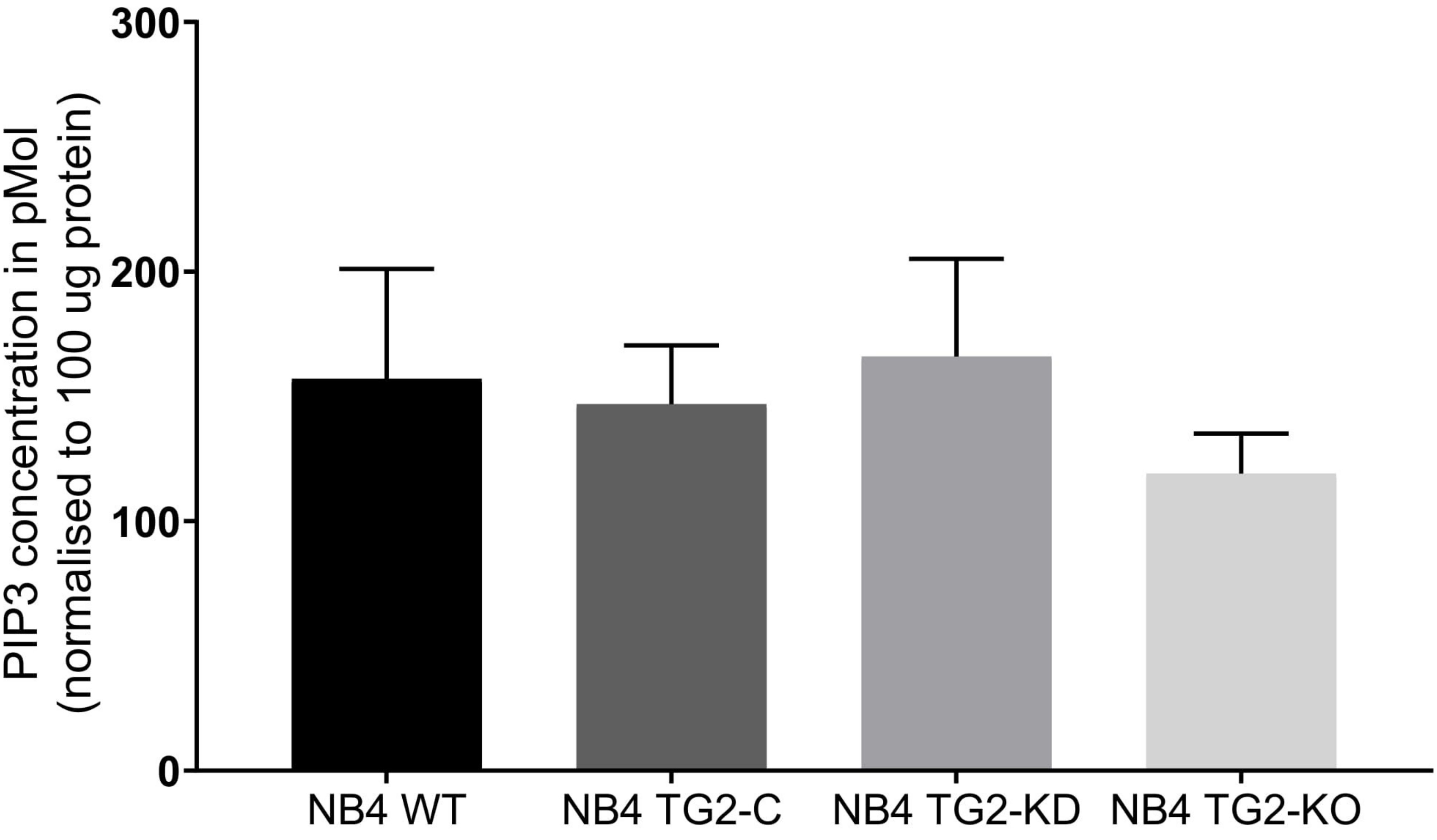

#### Supplemental Figures Legend

**Supplemental Figure 1. ATO-induced cell death.** (A) Cell number changes measured in KOVA Glasstic® Slide cell number counting chambers for NB4 WT, NB4 TG2-KD, and NB4 TG2-KO cells without treatment on the indicated days (n=9). The percentage of cell death is represented as mean % ± SD (n=9). (**B-C**) Cell number changes measured in KOVA Glasstic® Slide cell number counting chambers for NB4 WT, NB4 TG2-KD, and NB4 TG2-KO cells treated with 0.5 µM or 2.0 µM ATO on the indicated days (n=9). (**E-F**) FACS analysis of Annexin-V and PI-stained NB4 cell lines following ATO treatments. The percentage of cell death is represented as mean % ± SD (n=9). Measurements were conducted in triplicate; values were validated by Flowing software 2.5.1. **(G-I)** Detection of cell death in ATRA +ATO treated NB4 cells after 5 days. FACS analysis of Annexin-V and PI stained NB4 cell lines following ATRA+ATO treatment at day 5. The percentage of cell death is calculated by using the quadrant method. (G) The first row represents the control measurements by which the quadrant was determined. (H-I) Graphs represent the ATRA + ATO treated NB4 WT and NB4 TG2-KO cells raw data at day 5. Measurements were conducted in triplicates; values were validated by Flowing software 2.5.1.

**Supplemental Figure 2. Expression of "non-active" leukocyte β2 integrin CD11b/CD18 and CD11c/CD18.** (**A-C**) mRNA expression of differentiation markers CD11b in untreated or ATO treated NB4 cells (n=3). (**D-F**) Flow cytometry analysis of cell surface expression of non-activated differentiation marker CD11b (n=3). (**G-I**) mRNA expression of differentiation markers CD11c in untreated or ATO treated NB4 cells (n=3). (**J-L**) Flow cytometry analysis of cell surface expression of non-activated differentiation marker CD11c/CD18 (n=3). (**M-O**) Flow cytometry analysis of cell surface expression of non-activated differentiation marker CD11b/CD18 upon ATRA + ATO treatment (n=3). (**P-R**) Flow cytometry analysis of cell surface expression of non-activated differentiation marker CD11c/CD18 upon ATRA + ATO treatment (n=3). Measurements were conducted in triplicate; values were validated by Flowing software 2.5.1.

**Supplement Figure 3. CBX inhibits the FOXO3 pathway.** (**A**) NB4 WT cells containing a FOXO3A luciferase reporter element were treated with ATRA and ATO and measured using a luminescence-based method in triplicate, with values reported as relative light units (RLU) (n=3). Values were normalized to 100 µg cell lysate protein. (B) NB4 WT cells containing a FOXO3A luciferase reporter element were treated with ATRA+ATO with or without the FOXO3A inhibitor CBX at different concentrations. Graph represents the percentage of cell death measured by Annexin-V and PI staining upon FOXO3A inhibition by CBX. Statistical analysis was conducted by two-way ANOVA (Bonferroni post hoc test; *P<0.05, **P<0.01 and ***P<0.001, ****P<0.0001). Asterisk show the significant differences in ATRA+ATO treated vs. ATRA+ATO+CBX treated cells.

**Supplement Figure 4. $PIP_3$ concentration in undifferentiated NB4 cells.** $PIP_3$ levels in membranes of untreated NB4 cell lines were quantified by ELISA (Echelon Inc.). The mean values ± SD are shown for $PIP_3$ levels from three independent experiments. Measurements were conducted in triplicate, and the values were normalized against the protein concentration of the samples.

#### Supplemental METHODS

##### Fluorescence-activated cell sorting (FACS)

Approximately 4-6 × $10^5$ NB4 treated cells were harvested and washed with pre-cooled 1X PBS, followed by centrifugation at 1016 RCF (g) for 3 min at 4°C. All the following steps were performed at 4°C. Pellets were washed and incubated with 2% BSA containing 1X PBS for 15 min and then centrifuged at 1016 RCF (g) for 3 min. The centrifugation was followed by a 2 h incubation in the dark with phycoerythrin (PE) and FITC or allophycocyanin (APC) labeled CD11c/CD11b antibodies in a 1:25 dilution ratio (R&D Systems MAB16991, Biolegend). For each treatment isotype control were used for normalizing. Incubation with the antibodies followed by repeated washing steps, and the labeled samples were measured by FACS (BD FACScalibur instrument, BD FACSAria™ III flow cytometer BD Biosciences, San Jose, CA). Dead cells were excluded from the analysis by forward scatter (FSC) and side scatter (SSC) gating methods. Data were validated by Flowing software version 2.0.4, normalized, and corrected to the isotype controls of each antibody/treatment.

##### Annexin-V labeling of NB4 cells and live/dead cell sorting

Approximately 1-2 × $10^6$ NB4 treated cells were harvested and washed with pre-cooled 1X PBS, followed by centrifugation at 55 RCF (g) for 3 min at 4°C. All the following steps were performed at 4°C. Cells were labeled with FITC-conjugated Annexin-V (Biolegend) for 15 min in the dark. The excess amount of dye was removed by centrifugation at 55 RCF (g) for 3 min. The pellet was resuspended in pre-cooled PBS. Cells were analyzed and sorted on a BD FACSAria™ III flow cytometer (BD Biosciences, San Jose, CA). Cells were lysed and prepared for Western blot analysis.

For the apoptotic feature evaluation experiments, the sorting method was that first the size and the granularity features of the NB4 cells were used, followed by the gated cell population filtering against the FITC-positive cells. FITC-positive cells were sorted out into a new 15 mL tube. With each cell line, a minimum of 1 × $10^5$ cells were sorted based on their FITC positivity.

##### Gene expression

Isolation of RNA and RT-PCR/RT-QPCR methods have been published previously[17,18]. For the real-time Q-PCR reaction, the following TaqMan probes (ABI, Applied Biosystems) were used: TG2, PTEN, PI3K-p85, PI3K-p110, human ALU, GAPDH (Thermo Fisher Scientific) (Table

1). The analysis was carried out using the Roche LightCycler® 480 II (© 2021 Roche Molecular Systems, Inc)

| OLIGO | Assay ID |
|---|---|
| TGM2 | Hs01096680_m1 |
| PTEN | Hs02621230_s1 |
| PI3K-p85 | Hs00388782_m1 |
| PI3K-p110 | Hs00898499_m1 |
| Human-ALU sequence | Forward primer TGGTGGCTCTCTCCTGTAAT |
| Human-ALU probe | TGAGGCAGGAGAATCGCTTGAACC FAM-MGB |
| Human-ALU sequence | Reverse primer GATCTCGGCTCACTGCAAC |
| GAPDH | Mm01328877_g1 |
| CYCLOPHILIN-D | Mm01328877_g1 |
| ACTIN | Hs01101944_s1 |

*Table 1. Oligos used in RT-QPCR*

**Western-blot analysis**

The pellet was lysed in lysis buffer (50 mM TRIS, 1mM EDTA, 0.1 % MEA, 0,5% Triton X-100, 1 mM PMSF) containing a protease inhibitor cocktail (Sigma-Aldrich) with a 1:100 dilution ratio and homogenized with 5–7 strokes with sonicator at 40% cycle intensity (Branson Sonifer, 450). After sonication, the lysed samples were centrifuged at maximum RCF at 4ºC for 15 min. The supernatant was collected for measurement of the protein concentration.

The protein concentration was measured with the Bradford assay at a wavelength of 595 nm (Synergy Multi-Mode Microplate Reader). Every sample was measured with 3 technical parallels, normalized using BSA standard (Sigma-Aldrich, Stock: 0.5 mg/mL). The protein samples were diluted up to 2 mg/mL concentration, mixed with equal volumes of 2×SDS denaturation-buffer (0.125 M Tris-HCl, pH 6.8, containing 4% SDS, 20% glycerol, 10% MEA, 0.02% bromophenol blue), and incubated at 99°C for 10 min.

Depending on the molecular weight, proteins were separated on 8–15% SDS-polyacrylamide gels and blotted onto a PVDF membrane (MERCK-Millipore) using wet and semi-dry blotting methods. The wet blotting method was applied in the case of the mTOR proteins. The membranes were blocked with 5% non-fat dry milk/5% BSA in Tris-buffered saline and Tween 20 (TTBS) for 1 h at RT. Primary antibodies were diluted in 0.5% milk/5% BSA in TTBS, with a dilution ratio of 1:1000–1:5000, incubated overnight at 4°C (Table 2). The membranes were washed three times with TTBS for 15 min at RT, incubated with horseradish peroxidaselabeled, affinity-purified secondary antibodies (Advansta) at a 1:10000-1:20000 dilution ratio for 1 h at RT, and then washed three times with TTBS for 15 min at RT. The targeted protein band was visualized using the ECL-Kit (Advansta). Quantification of the protein bands was performed using ImageJ software version 1.09.

| ANTIBODIES | DILUTION RATIO |
|---|---|
| p-PTEN (S380/T382/383) rabbit-anti human | 1:1000 |
| PTEN rabbit-anti human | 1:5000 |
| p-mTOR (S2448) rabbit-anti-human | 1:1000 |
| p-mTOR (S2481) rabbit-anti human | 1:1000 |
| mTOR rabbit-anti human | 1:5000 |
| p-AKT (T308) | 1:5000 |
| p-AKT (S473) | 1:5000 |
| AKT | 1:5000 |
| p-PI3K p85 (Y458)/p55 (Y199) rabbit-anti human | 1:5000 |
| PI3 kinase p85 (19H8) rabbit-anti human mAb | 1:1000 |
| PI3K p110 delta subunit | 1:1000 |
| tTG anti-polyclonal | 1:5000 |
| PI3K p110beta (c-8) rabbit polyclonal | 1:5000 |
| Caspase-3 (H-277) rabbit polyclonal | 1:3000 |
| p-FOXO1/FOXO3/FOXO4 (T24/T32) | 1:5000 |
| GAPDH anti-mouse-human FF26A/F9 clone | 1:5000 |
| FOXO3A W15111A clone | 1:1000 |
| PTEN 4C11A11 clone | 1:5000 |

*Table 2. Antibodies used for Western blotting*

#### $PIP_3$ isolation

After aspirating media from treated NB4 cell lines, 1 mL cold 0.5 M TCA were added. Samples were incubated on ice for 5 min and transferred into a 15 mL tube, followed by centrifugation at 13,000 RCF (g) for 5 min at 4°C. The pellets were washed with 1 mL 5% TCA/1 mM EDTA, vortexed, centrifuged again at 13,000 RCF (g) for 5 min at 4°C, and the supernatants were discarded. After the addition of 1 mL methanol:chloroform (2:1), the pellets were vortexed 3-4 times over 10 min at room temperature. Samples were centrifuged again at 13,000 RCF (g) for 5 min at 4°C, and 0.5 mL chloroform:methanol:12 N HCl (40:80:1) was added to the pellets. The samples were then incubated for 15 min with occasional vortexing. A 180 μL volume of chloroform was added, followed by 320 μL 0.1 N HCl. The samples were centrifuged at 13,000 RCF (g) for 5 min at 4°C. The organic (lower) phases were transferred to a new 1.5 mL tube, followed by the addition of 30 μL 0.1M ammonium hydroxide in methanol. The neutralized organic phase was then dried in a vacuum dryer.

**Human CD11c/CD11b/Annexin-V positive cell labeling from mouse blood**

Blood was collected from the hearts of mice at day 14 and treated with sodium citrate (8% v/v) to avoid coagulation. Blood samples were lysed with lysis buffer (BD PHarmn LyseTM) following the kit manufacturer's protocol. Lysed samples were centrifuged at 1016 RCF (g) for 3 min. The centrifugation was followed by a 2 h incubation in the dark with the PE and FITC or APS labelled CD11c/CD11b antibody in a 1:25 dilution ratio (R&D Systems). Samples were labeled with F4/80 antibody (Biolegend) in a 1:100 dilution ratio for mice macrophages for 15 min in the dark at 4°C. In the last step, after centrifugation, the supernatant was discarded. The cells were resuspended in Annexin-V binding buffer and labeled with APC conjugated Annexin-V (Biolegend) for 15 min at 4°C in the dark, followed by measurement with a BD FACSAria™ III flow cytometer.

Cells were first gated based on their size and granularity (FSC and SSC). From the total cell population, F4/80 positive mouse cells were excluded, and the human NB4 cell population was gated out. CD11 positive cells were gated out from the total human cell population in the next step. From the gated CD11 positive human NB4 cells, based on APC labeling, the Annexin-V positive cells were filtered out, resulting in a CD11 and Annexin-V positive cell population. FACS data were validated by Flowing software 2.0.5 version normalized and corrected to the respective isotype controls.

**Densitometric analysis**

Integrated optical densitometry was performed using ImageJ software version 1.09. Values were normalized to the loading control of each blot/sample/lane. Blot pictures were changed into RGB 32-bit mode, and based on the gray-scale format, the analysis was performed using fixed rectangular selection.

***In vivo* mouse experiment**

All experiments were performed according to the guidelines of the Institute for Laboratory Animal Research, University of Debrecen, Faculty of Medicine, and were approved by the national and institutional ethics committee for laboratory animals used in experimental research (Project ID:4/2020/DEMAB).

NB4 WT and NB4 TG2-KO cells ($1\times10^7$) were washed with sterile phosphate buffered saline (PBS) and injected into the retro-orbital region of 8–9-week-old NOD/SCID (CB17/lcr-Prkdc$^{scid}$, JANVIER LABS) female mice under sterile conditions.

Experiments were carried out with four groups of treatments:

1. Control [DMSO:Ethanol (1:1)]
2. All-trans retinoic acid (ATRA) at a 1 μM final concentration + Dexamethasone (DEX) 50 μg
3. Arsenic trioxide (ATO) at a 2.0 μM final concentration
4. ATRA 1 μM + ATO 2.0 μM

Treatments were performed every two days in anesthetized mice at 1.0 mg/kg ATRA + 50 μg DEX, 0.75 mg/kg ATO, or in a combination as ATRA + ATO intraperitoneally for 14 days. Blood samples were harvested from the tail veins of the mice at the beginning of treatment and again at the termination on day 14. A 27.5-gauge or 0.5-gauge insulin needle/syringe (Terumo U-100, Terumo Medical Corporation, Elkton, MD) was used for the injections of the treatment i.p. As an experimental control, a DMSO:Ethanol solution was administered i.p. (the ATRA powder stock solution was prepared in DMSO-ethanol. While performing the i.p. injection of the animals, the DMSO (v/v%) values were kept under 2% to avoid any cytotoxic effects. At the end of treatment at day 14, mice were anesthetized with isoflurane. Circulating human NB4 and mouse cells were analyzed from total cardiac blood samples with flow cytometry and distinguished by human and mouse neutrophil surface-expressed marker proteins.

**Labeling of mouse blood with human CD11c/CD11b/Annexin-V positive cells**

Blood was collected from the hearts of 14-day-treated mice into 8% sodium citrate-treated tubes, lysed in lysis buffer (BD PHarmn Lyse™), centrifuged at 55 RCF (g) for 3 min, and then incubated for 2 h in the dark with PE-labeled CD11c and FITC-labeled CD11b antibodies (1:25 dilution; R&D Systems). Macrophages in the samples were then labeled with F4/80 antibody (1:100 dilution; Biolegend) for 15 min in the dark at 4°C. The cells were pelleted by centrifugation, resuspended in Annexin-V binding buffer, labeled with APC-conjugated Annexin-V (Biolegend) for 15 min at 4°C in the dark, and then analyzed with a BD FACSAria™ III flow cytometer. F4/80 positive mouse cells were excluded, and the human NB4 cell population was gated out. CD11 positive cells were then gated out from the total human cell population, and APC labeling was used to filter out the Annexin-V positive cells, resulting in a CD11 and Annexin-V positive cell population. The FACS data were validated with Flowing software version 2.0.5 and normalized and corrected with appropriate isotype controls.
